# An Innovative Setup for High-Throughput Respirometry of Small Aquatic Animals

**DOI:** 10.1101/2020.01.20.912469

**Authors:** M.K. Drown, A.N. DeLiberto, D.L. Crawford, M.F. Oleksiak

## Abstract

Metabolic rate is often measured as a phenotype in evolutionary genetics studies because it impacts organismal fitness, is repeatable and heritable, and is responsive to numerous environmental variables. Despite a wide body of literature about metabolic rates, key questions remain unanswered: 1) why do individuals from the same population exhibit up to three fold differences in metabolic rate, 2) how does metabolic rate change during an individual’s lifetime, and 3) what metabolic rate is advantageous in a specific environment? Current low throughput approaches to measure metabolic rate make it difficult to answer these and other relevant ecological and evolutionary questions that require a much larger sample size. Here we describe a scalable high-throughput intermittent flow respirometer (HIFR) design and use it to measure the metabolic rates of 20 aquatic animals simultaneously while reducing equipment costs and time by more than 50%.

## Introduction

Metabolic rate is often measured as a phenotype in evolutionary genetics studies because it is known to impact organismal fitness, is repeatable and heritable, and is affected by a variety of environmental variables (1–5). The relationship between metabolic rate and a variable of interest, such as temperature, oxygen availability, or toxicant exposure, has been investigated frequently, which has led to a rich literature on metabolic rates in many species (7–11). Despite this wide body of literature, key questions about metabolic rates remain unanswered including 1) why do individuals from the same population exhibit up to three fold differences in metabolic rate under similar acclimation conditions and activity levels, 2) how does metabolic rate change during an individual’s lifetime, and 3) what metabolic rate is advantageous in a specific environment (7)?

Flow through respirometry, intermittent-flow respirometry (IFR), and closed respirometry are techniques used to measure metabolic rates in terrestrial and aquatic organisms. Flow through respirometry is achieved by measuring the amount of oxygen entering and leaving a chamber relative to the flow rate of air or water through the chamber (12). In IFR the respirometer cycles between open and closed periods. During open periods the chamber is flushed to remove waste and oxygen is replenished and during closed periods the animal is using oxygen sealed in the chamber (Fig. 1) (12, 13). Closed respirometry places an organism in a sealed chamber of known volume and measures oxygen or carbon dioxide partial pressures at multiple time points throughout the trial. The sealed chamber during closed respirometry may result in the accumulation of nitrogenous waste and carbon dioxide, which can increase stress, and may cause loss of equilibrium (LOE) in aquatic organisms (14).

**Figure 1:**
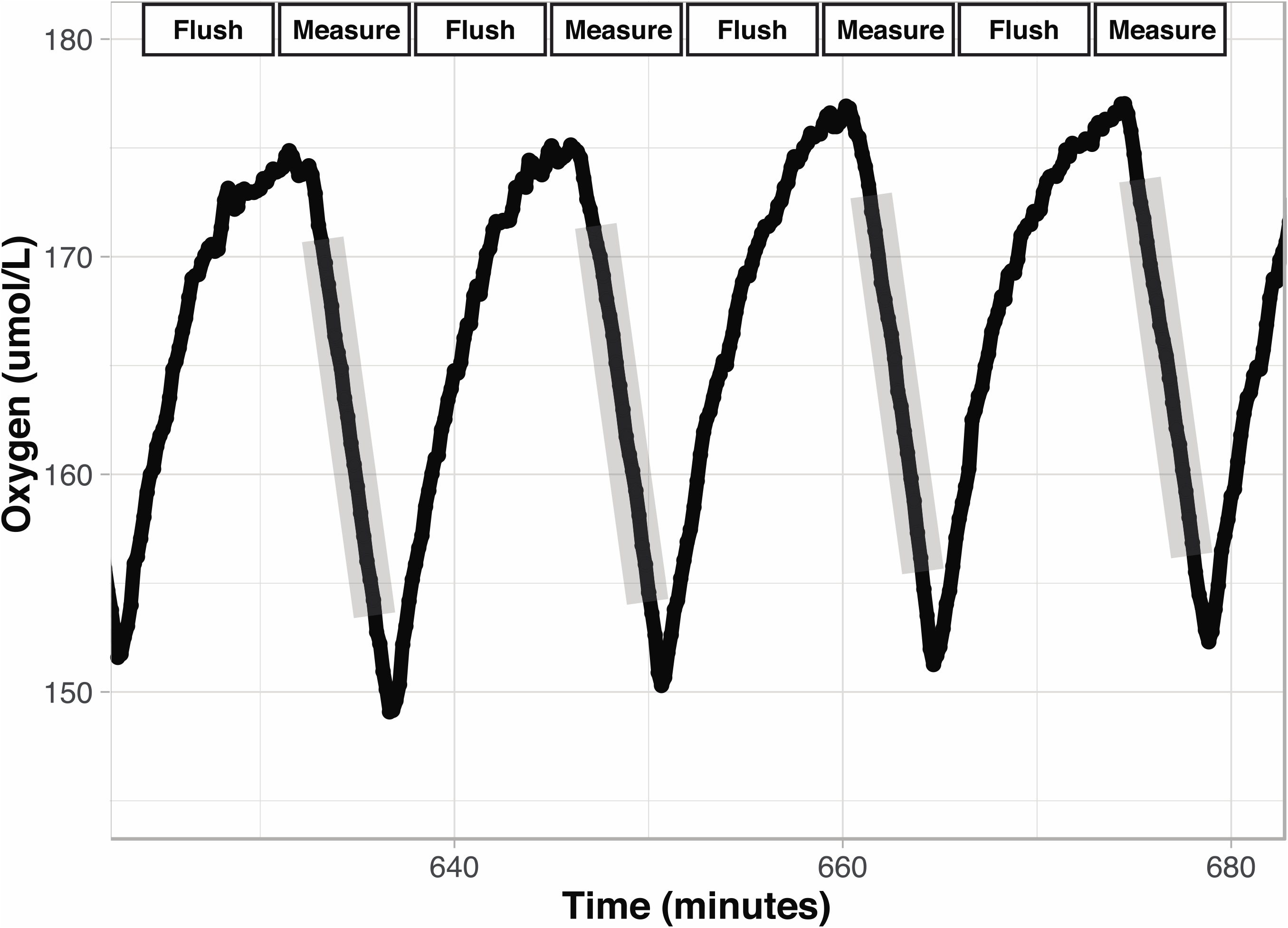
Intermittent Flow Respirometry. Oxygen concentration over time in a chamber during intermittent flow respirometry. There are two period types: Measurement and Flush. Measurement periods (gray shading) occur when the chamber is sealed, and the decrease in oxygen concentration reflects oxygen consumption by the organism. The slopes of the lines (oxygen *vs*. time) during the measurement periods are used to calculate metabolic rate. Flush periods are when the rapid increase in oxygen occurs as fully oxygenated water is pump into the chamber. Data displayed are from the setup described here.

Flow-through respirometry and IFR methods may be applied to measure standard, resting, or maximum metabolic rate. In contrast, closed respirometry uses a single closed period and yields an average metabolic rate based on oxygen consumption at multiple time points as oxygen declines in the chamber (15). Standard metabolic rate (SMR) is measured when an animal is at rest and in a post-absorptive state (i.e., fasting). Routine metabolic rate (RMR) is similar to SMR but includes spontaneous activity in animals that do not have a motionless rest cycle (16). Maximum metabolic rate (MMR) is the highest maintainable metabolic rate an individual can achieve (13).

To measure metabolic rate, swim tunnels or respirometers can be purchased. Both are widely available with a variety of oxygen sensing technologies and software packages. Typically, these swim tunnels or respirometers are designed to house one organism at a time for several hours or days in order to achieve a precise measure of metabolic rate, and they are often expensive to purchase as a complete measurement system (~$20,000). Some companies additionally offer high-throughput versions (up to 8 chambers) for small animals; however, the cost remains high (>$2000 per chamber when including software and oxygen sensing technology, ex: Loligo complete mini chamber system) with variable cost depending on the size of the chambers desired. While some authors have indicated that they have the capacity to measure 8 or more individuals simultaneously and report measuring dozens of individuals (17, 18), details of procedures and methods used as well as cost effectiveness are not publicly available. These restrictions make it challenging to measure metabolic rate rapidly for a large number of individuals without introducing time bias as the first and last individual may be measured weeks or months apart depending on sample size. With little known about the way metabolic rate changes within an individual’s lifetime, it is difficult to know how much variation among individuals is due to time *versus* physiological differences in experiments entailing weeks to months between the first and last individual measurements (7). The limitation on throughput additionally prevents questions relevant to ecology and evolution from being answered as these questions often require a much larger sample size than can feasibly be measured with current available methodologies. Given the high interindividual variation in metabolic rate, characterizing ten or twenty individuals at a given life stage or under specific treatment conditions may not capture the scope and shape of the physiological response to various stressors within a population or species and would inhibit the discovery of broad patterns across taxa (2, 7). Additionally, due to the plasticity of metabolic rates in some species, measuring metabolic rate at one time point in one environment may not reflect an ecologically relevant trait. Thus, our ability to understand variation in metabolic rate will be limited until we are able to reasonably measure larger sample sizes for species or populations of interest or to obtain repeated measures of the same individuals across various timepoints and in various environments. Here we describe a scalable high-throughput intermittent flow respirometer (HIFR) design and use it to measure the metabolic rate of 20 aquatic animals simultaneously, which may allow us to achieve the large sample sizes needed to answer these complex questions.

## Materials and Methods

The custom HIFR system is a large water bath with a PVC rack that holds 20 glass chambers. Each chamber has tubing and pumps to flush the chamber and re-circulate water that passes by an oxygen sensor (Fig. 2).

**Figure 2:**
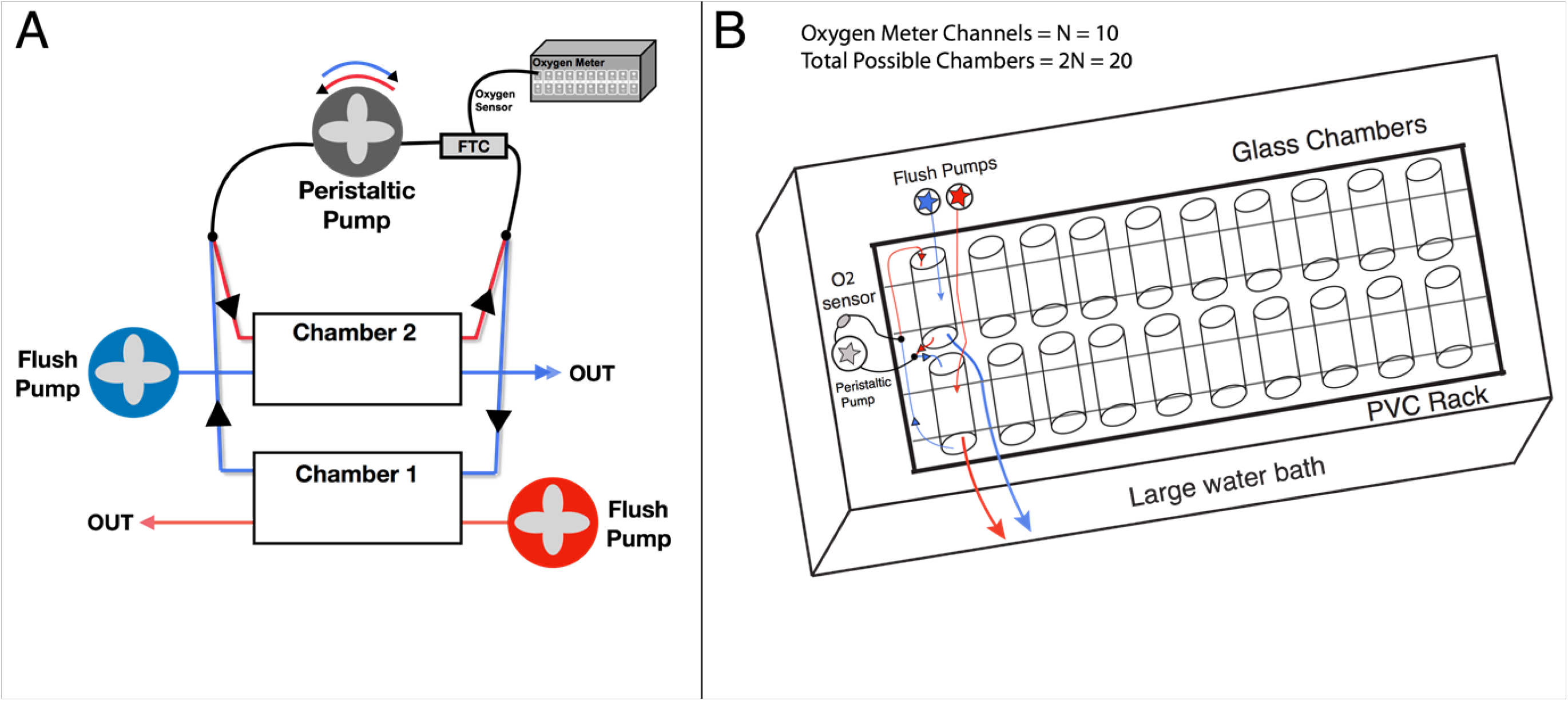
Pumping Circuits. **A)** Pairs of chambers in the high-throughput intermittent flow respirometer. Circuit 1 (red), circuit 2 (blue). One-way values (black arrows) control flow direction. By changing the polarity of the peristaltic pump motor, the peristaltic pump direction changes. **B)** Overall schematic of HIFR. The basic design is a PVC rack that holds and secures glass chambers with their rubber stoppers, which is placed in a large water bath. Each chamber is connected to flush pumps and re-circulating pumps with oxygen sensors. Throughput is limited by the number of channels on the oxygen meter (N) with this design able to measure 2N individuals simultaneously.

The large water bath (1.2 m long, 1.1 m wide, 0.3 m deep) was constructed out of 0.635 cm thick plexiglass and sealed with plastic weld and silicone glue to prevent leaking. A PVC rack with twenty slots separated by small PVC pieces was placed in the water bath, and a 0.300 L glass chamber was placed in each slot and sealed with two rubber stoppers. Each rubber stopper had two 0.635 cm stainless steel tubes to attach flexible tubing connected to pumps. Glass chambers were then paired and attached to one peristaltic pump (60 mL/minute) and two separate flush pumps with flow directed to 10 chambers each (300 mL/minute per chamber). A flow-through-cell with an oxygen sensor with a fiber optic cable (FTC) was placed in line with the peristaltic pump. The fiber optic oxygen sensor was attached to a 10-channel oxygen meter (PreSens Precision Sensing, Regensburg Germany). A separate PT-100 temperature probe was placed in the large water bath (PreSens Precision Sensing, Regensburg Germany). PreSens Measurement Studio 2.0 software was used to record oxygen over time as the peristaltic pump recirculated water through the FTC and past the oxygen sensor then back to the chamber.

An Arduino Uno with a 5V relay and a set of double pole double throw relays was used to control the direction the peristaltic pump turned and the power to the flush pumps. One-way valves were used to control the flow path to and from the peristaltic pump such that when the pump rotated clockwise, it would draw water from one chamber past the FTC, and when the pump rotated counter-clockwise, it would draw water from a second chamber (Fig. 2). Thus, all the odd numbered chambers be measured while the even numbered chambers were being flushed and *vice versa*. The one-way values also prevented the back-flux of water so that water was not mixed between sets of chambers. In this way the ten peristaltic pumps with FTCs were used to measure oxygen levels in 10 of the 20 chambers at one time and would oscillate between sets of 10 by changing the polarity of the peristaltic pump motor. Altering flow between the ten FTC and the flush circuit allowed the 10-channel oxygen meter to measure the twenty respirometers housed in the temperature-controlled water bath. A detailed list of materials and costs for building a HIFR can be found in Table S1, and a schematic of the electric circuit used to control and power pumps is depicted in Figure S1.

### Animal care and use

*Fundulus heteroclitus* are a small estuarine fish often used to address questions in physiology and genetics because they are known to be highly plastic and adaptable to changing environments. Found from New Brunswick, Canada to northern Florida along the East Coast of the United States, *F. heteroclitus* live along a thermal cline of at least 14°C and additionally experience variation in temperature, salinity, water depth, and dissolved oxygen with daily tidal cycles and seasonal weather changes (19, 20).

*F. heteroclitus* were caught in live traps along the east coast of the United States in New Jersey and transported live to the University of Miami where they were housed according to the institutional animal care and use committee guidelines (Animal Use Protocol No: 16-127-adm04). *F. heteroclitus* were collected on public lands and do not require a permit for non-profit use. Fish were common gardened at 20°C 15ppt for greater than 6 weeks on a summer light cycle (14 hours daylight, 10 hours dark), overwintered at 10°C and 15ppt (5 hours daylight 19 hours dark) for 4 weeks, and then acclimated to 28°C and 15ppt for at least four weeks on a summer light cycle (14 hours daylight, 10 hours dark). Fish were fed pelleted food to saturation once daily and fasted for 24 hours prior to metabolic rate determination.

To identify individuals, all fish had unique visual implant elastomer (VIE) tags. Metabolic rates were measures after at least four weeks of acclimation to 28°C.

### Metabolic rate calculation

Individuals were measured overnight where they were left undisturbed for at least 14 hours. Fish were identified then immediately placed in a chamber between 16:30h and 17:30h and the first replicate measurement period began after midnight, allowing a minimum of 6.5 hours of acclimation to the chamber. Each measurement period lasted 6 minutes followed by a 6-minute flush period to prevent oxygen levels from dropping below 80% in any given chamber and to fully replenish oxygen levels in the chamber between replicate measurements. The water bath housing the twenty chambers was continuously recirculating with an aquarium system containing a biofilter of nitrogen fixing bacteria to reduce ammonium load and a heating unit that kept the water bath at the desired temperature (±1°C).

After each night, data files were exported from PreSens measurement studio and analysis was done using R (version 1.1.383). From each 6-minute measurement, the first and last 1 minutes were excluded as a buffer between the flush pump turning off and the measurement period starting. An R-Markdown script detailing the processing of raw data files is available on github (https://github.com/mxd1288/FunHe_Genomics/blob/master/Raw_Metabolic_Rate_Pipeline.Rmd).

The slope of oxygen levels over time was extracted using a linear model for each replicate measurement period, and MO_2_ in μmol O_2_l^−1^ was calculated using the equation y=KV then converted to mg O_2_l^−1^ for comparability, where y=MO_2_ (μmol O_2_min^−1^), K=slope (μmol min^−1^), V= volume of the respirometer (including tubing) minus volume of the organism (liters) (13). Any data collected while the lights were on in the room (before 23:00h or after 06:30h) or a slope with an R^2^ value less than 0.9 were excluded from the analysis. Between midnight and 06:30h at least 25 measurement periods were completed for each individual, of those at least 20 were used for analysis after exclusion based on R^2^ value. The lower 10^th^ percentile values from the cumulative frequency distribution of all replicates from that individual were used to estimate standard metabolic rate (SMR). Using the lower 10^th^ percentile value from the cumulative frequency distribution did not average the lowest two metabolic rate measures. One value for each individual that lay on the continuous cumulative frequency distribution at the 10^th^ percentile was selected to represent each individual. This lower 10^th^ percentile value captures the time period when the fish were most at rest during measurement and excludes the lowest tail of the data distribution, which may be sensitive to outliers (16, 21, 22). (Fig. 3).

**Figure 3:**
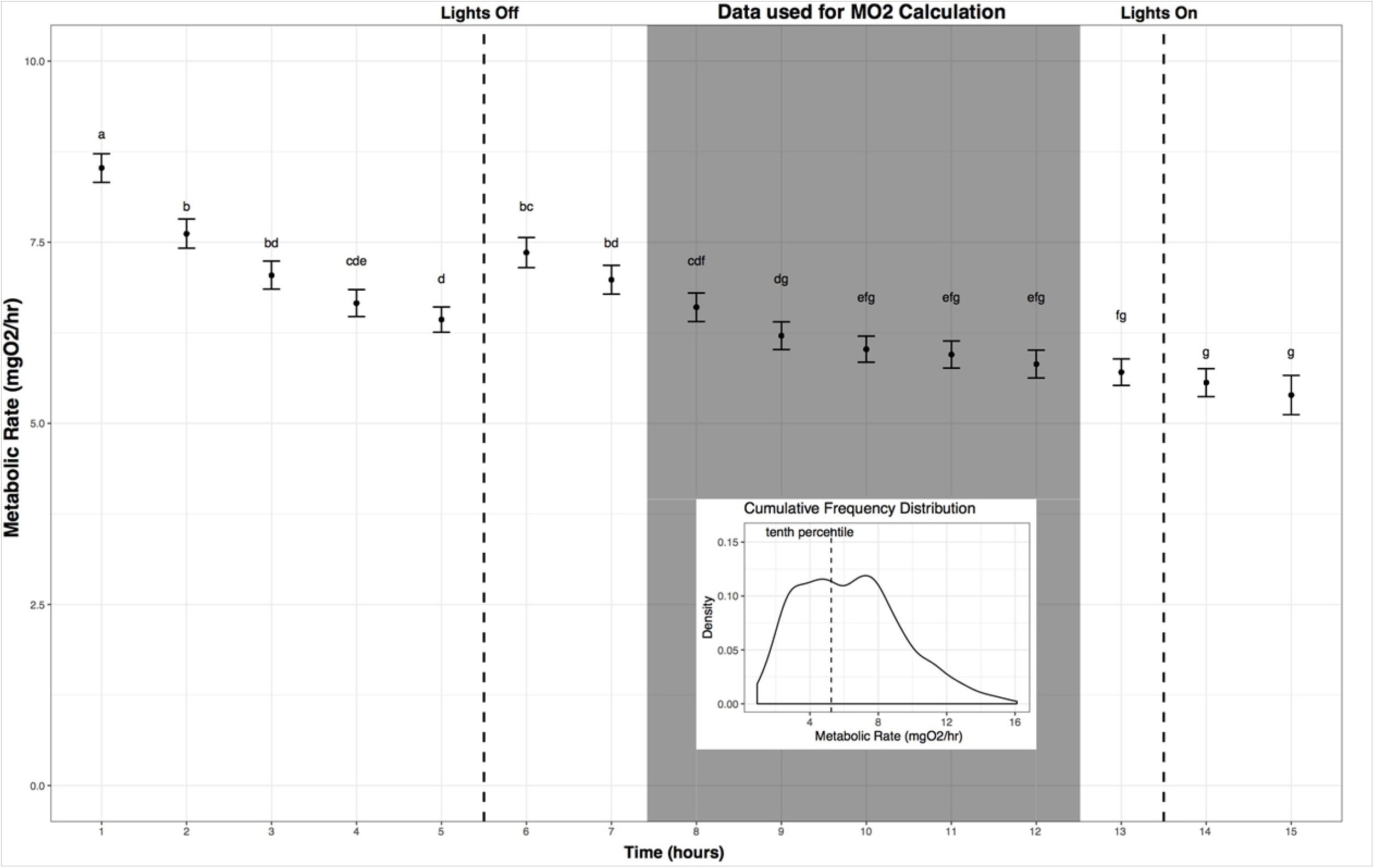
Metabolic Rate Measurement Over Time. Metabolic rate *versus* time since fish were added to the chambers (mean and standard error across all individuals on an hourly basis). Fish reached a resting state in the chamber between 3 and 4 hours when left undisturbed. Replicates used in calculating metabolic rate (MO_2_) are indicated (shaded box). Letters indicate significant differences among time points (ANOVA, α=0.05). **Inset:** The lower 10^th^ percentile values from the cumulative frequency distribution of this subset of replicates were used to estimate standard metabolic rate for each individual.

### Body mass correction

To compare metabolic rates among individuals that vary in size, metabolic rate must be corrected for body mass. Fish were weighed to the nearest 0.1g the day of metabolic rate measurement. After calculating SMR the residuals of the model metabolic rate (log transformed) *vs*. body mass (log transformed) were used as the body mass corrected SMR (23).

### Background respiration

In order to correct for oxygen used by bacteria and other microorganisms in the HIFR, blank runs were completed in between each use of the HIFR and average background respiration subtracted from the MO_2_ of each fish (Eq. 2).

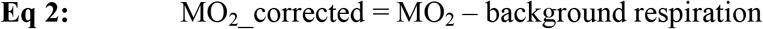

Where MO_2_ is the minimum metabolic rate of each fish as previously described and background respiration was a chamber specific value calculated by averaging the oxygen consumption over time in each empty chamber across three replicate blank runs. An empty chamber was additionally run in parallel each night and the background respiration did not change over the course of the night validating the decision to not use a time corrected value of background respiration.

## Results and Discussion

### System design and testing

Water at 28°C (±1°C) and 15ppts was used to fill the custom water bath and recirculated with an aquarium system to maintain temperature and reduce ammonium load. To validate that the flush period was long enough to fully replenish oxygen empty chambers were filled with water at a low oxygen concentration (~60% a.s.), achieved by bubbling in nitrogen, and flushed for over 8 minutes. Between 4 and 5 minutes after turning on the flush pump the oxygen level in the chamber exceeded 99% a.s. (Fig. S2). Using the equation for steady-state transformation shows that with a flush rate of 300mL/min, and a chamber that is 300mL in volume, water should reach 99% replacement after 4.61 minutes (t(99%) = −ln(1 − 0.99) × 300 mL / 300 mL/min = 4.61 min, (6)).

To test system leak one chamber in a pair was filled with water at oxygen concentration equal to ~50% a.s. and the other chamber in the pair was sealed with the flush pump running. A model of oxygen *versus* log10(time) from 75% a.s. to 89% a.s. (maximum oxygen reached) was used to derive an equation that can be used to predict the amount of leak at a specific time point: slope (% a.s. per minute) = 7.4916/time (minutes)(Fig. S3). Note that this equation can be applied from the time when 75% a.s. was reached (16.5 minutes) and extrapolated to determine time to reach 100% a.s. (144.5 minutes) but not used to predict oxygen concentration at previous time points not included in the model. Leak did not exceed 0.5% a.s. per minute from 80 to 89% a.s. and decreased as the oxygen concentration in the chamber increased. At 85% a.s. or higher leak would not exceed 0.14% a.s per minute. Thus from 100% a.s. down to 85% a.s. leak would be negligibly low (below 6% of typical MO_2_). Below 85% a.s. leak would increase, however, the portion of the slope used to calculate MO_2_, as described above (see metabolic rate calculation), would exclude the time period where oxygen levels would drop this low limiting the overall system leak. Leak could be reduced further by using material less permeable to oxygen to seal the chambers, although the cost of these materials may be higher (24).

### Repeatability

A random set of 19 fish was measured in the HIFR, each in three different chambers over the course of one week (Monday, Wednesday, and Friday night). Log SMR was regressed against log body mass (y=2.66 + 1.08x, R^2^=0.59, N=57, Fig. 4A), and a body mass correction was calculated as described above. The mean coefficient of variation (CV) within an individual was 18.03% (Fig. 4B). SMR is repeatable (Fig. 4C), and the variance for each individual for three SMR measured in three different chambers is much smaller than the variance among individuals (ratio of variance in group means/mean of within individual variance = 74.54:1). To measure repeatability (R) directly: 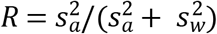 where s^2^_a_ equals the difference in the mean sum of squares among and within individuals divided by the number of measures per individual and s^2^_w_ equals the mean sum of squares within individuals (25, 26). The mean sum of squares among and within individuals can be taken from ANOVA (within = 0.0108, among = 0.8050) and using 3 measures per individual yields a repeatability of the tenth percentile value of metabolic rate, used here to represent SMR, of 0.96.

**Figure 4:**
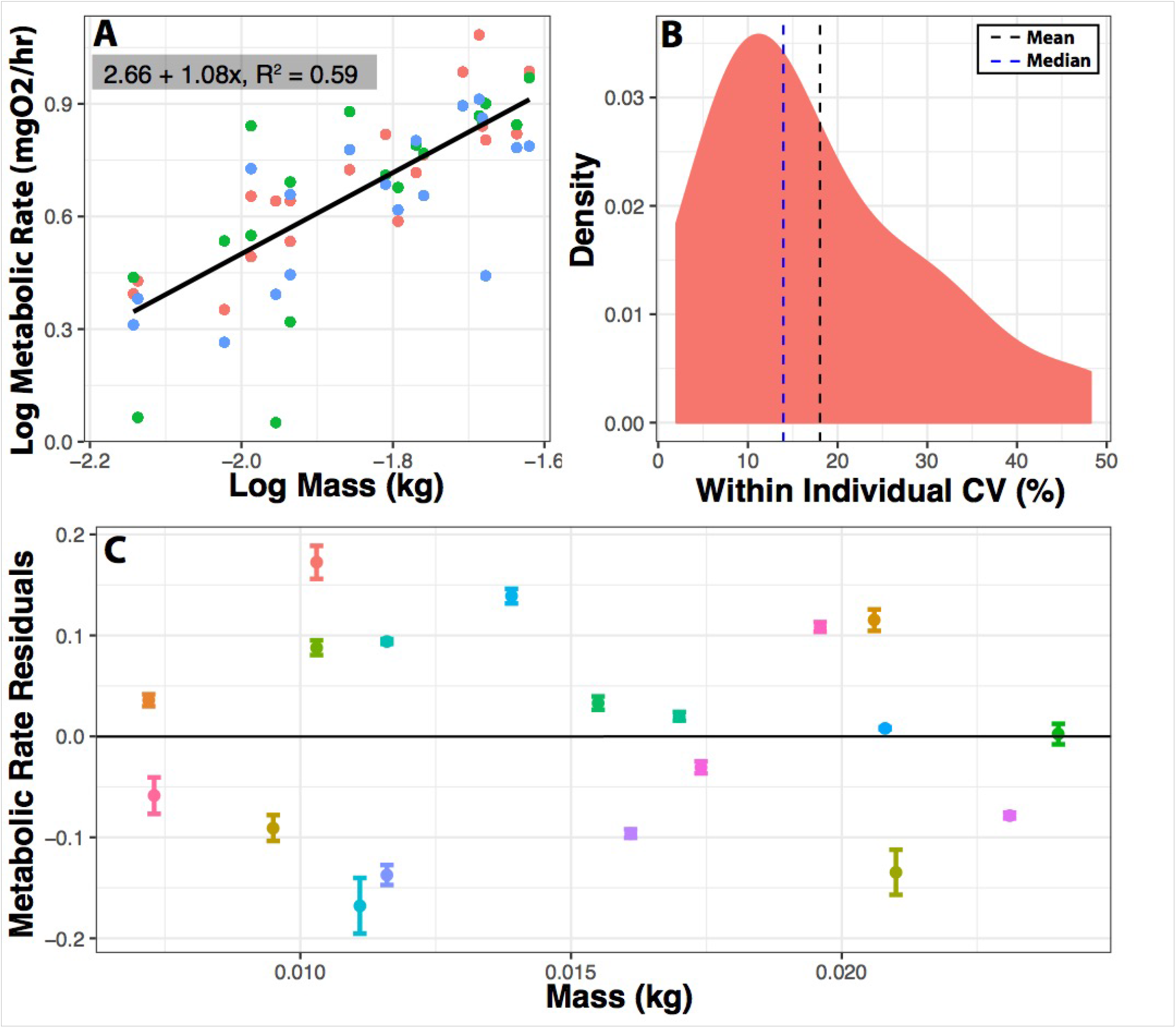
Repeatability of Metabolic Measurements. Metabolic rate was measured three times for 19 individuals in three different chambers within one week. **A)** Log metabolic rate (mgO_2_/hr) vs. log body mass regression. Values are corrected for background respiration. **B)** Distribution of coefficient of variation (CV) in minimum metabolic rate within an individual. CV = 100*(standard deviation/mean). **C)** Mean and standard error for SMR residuals among 19 individuals. Means are residuals from log-log body mass regression. Thus, positive values indicate that an individual had a higher than expected metabolic rate based on mass, and negative values indicate that an individual had lower than expected metabolic rate based on mass. Ratio of variance between to variance within = 74.82:1. Repeatability = 0.96.

Metabolic rates measured in HIFR are comparable with values from previously reported metabolic rate values for *F. heteroclitus* (±5%) and other teleost fish (±40%) further validating the methods described here (16, 27, 28). To determine this, the metabolic rate reported for *F. heteroclitus* or other species acclimated to various temperatures was used to interpolate metabolic rate for an 8-gram individual at 28°C. Differences between values reported from our HIFR and other studies using *F. heteroclitus* may also be due to the type of metabolic rate (i.e. SMR, RMR, MMR) being measured and the environmental parameters (acclimation *vs*. acute temperature or hypoxia exposure, maximum *vs*. standard metabolic rate, *etc*.). The 18.03% CV within individuals for SMR is similar to previous studies that reported 12-14% CV in SMR, MMR, and aerobic scope of brown trout (29).

Depending on the oxygen sensing technology and software, a single respirometer (including the cost of oxygen sensing technology and software) may cost between $2,000 and $4,000, a significant investment especially considering that they are designed to measure one individual at a time, which may take hours or days. Some companies additionally offer high-throughput versions (up to 8 chambers) for small animals; however, the cost remains high (>$2000 per chamber). The HIFR presented here was assembled using basic materials and a moderately priced oxygen meter and oxygen sensors. Including the cost of the meter, sensors, and materials, HIFR costs $855.50 per chamber to assemble, a 57% reduction in cost per respirometer compared to purchasing a Loligo high-throughput system. Additionally, the HIFR can simultaneously run up to 20 respirometers at once, greatly reducing the total time needed to achieve a large sample size, which holds value far beyond monetary savings. For example, within a one-week period at least 100 individuals could be run under the same experimental conditions introducing little variation due to time and requiring only 5 nights of respirometry set up with daily background respiration measures. The flexibility of the HIFR offers the additional advantage of allowing organisms of various sizes to be measured. By changing the size of the glass chambers and altering the flow rate of peristaltic pumps and flush pumps by changing the tubing size the system can easily be adapted to fit the desired organism. This further decreases costs for groups who may wish to measure a single species at various ages and stages of life or different species that may vastly differ in size (30).

Costs could be cut further by using less expensive peristaltic pumps or a different water bath than described here. However, the lifespan of a given pump varies greatly depending on the quality of the motor and the tubing. Several peristaltic pumps ranging from $3 to $50 were tested to determine the appropriate tubing material and motor design that could withstand frequent long-term use and alternation of motor polarity without rapidly burning out. Generally, it is recommended to use a peristaltic pump that has a brushed motor and tygon tubing and to determine the tubing size based on the desired flow rate. While there are large peristaltic pumps available it should be noted that depending on chamber size this may not provide enough mixing to prevent the stratification of water in the chamber (31). The addition of a closed loop mixing pump could mitigate this problem and provide adequate mixing, although this has not been tested here. Including the mixing pump would increase the total setup cost and without it the size of the chambers (and organisms) that can be measured with this system will be limited to those that can be adequately mixed with only a peristaltic pump.

The plexiglass tank served as a water bath for the chambers and could be replaced with a cheaper alternative as long as it could hold the appropriate volume of water needed to maintain a stable temperature and prevent the buildup of nitrogenous waste over the course of the run. It would also need to be large enough to hold the desired number of chambers of a specific size. In general, the respirometer volume should be 20 to 50 times larger than the organism to achieve a measurable decrease in oxygen over a reasonable period of time (several minutes) (6, 12). If the respirometer volume does not fit within this ratio for a given organism the measurement period length can be adjusted to allow for the appropriate drop in oxygen (above 80% O_2_ saturation) as long as routine movements are not inhibited by chamber size (6, 12). If variation in body mass of individuals is large, adjusting the measurement period to prevent low O_2_ levels for larger individuals may mean that smaller individuals do not have a large enough O_2_ decrease to get a reliable slope measurement. Using the HIFR design it is possible to use chambers of various sizes within a single run as long as pump flow rates were adjusted (tubing sizes) to accommodate this change. This allows for added flexibility of running different sized respirometry chambers simultaneously.

The throughput of this design is limited by the number of channels available on the oxygen meter. Any flow through oxygen sensing cells can be interchanged for the ones used here; however, the oxygen meter needs N channels (one per sensor) to allow measurement of 2N individuals at once. If an oxygen meter were available with 20 channels, for example, it is feasible that this design could be scaled to measure 40 individuals over the course of one night. An oxygen meter with fewer channels could be used to design a similar HIFR with fewer individual respirometers. Additionally, a HIFR could be built to measure ten individuals with ten channels by eliminating the double pole double throw relays and using an Arduino to turn the flush pump on and off.

Due to the size of the water bath and the available equipment, the most practical solution to maintaining a constant temperature in the water bath was to recirculate the water through a temperature-controlled aquaria system. This made it possible to pump the HIFR water bath into the same system the fish had been housed in prior to measuring metabolic rate so the temperature along with pH and salinity were not variable between the HIFR and the acclimation conditions (32).

It should be noted that the HIFR was built by an early career biology graduate student with little prior knowledge of electrical engineering or plumbing. The easy to learn techniques used make this methodology highly accessible.

The ability to precisely measure metabolic rates in a high-throughput manner without significantly increasing the necessary effort has application for physiologists, ecologist, geneticists, and comparative biologists alike. This method reduced the total system cost from ~$2,000 per respirometer to ~$900 per respirometer including the cost of the FTC, oxygen sensor, and oxygen meter. The HIFR also greatly reduced the effort needed to measure metabolic rate in a large sample size making it possible to answer questions relevant to ecological and evolutionary biology.

## Acknowledgements

Thanks to Dr. Chris Langdon for his advice and assistance in construction and to Moritz Ehrlich for constructive discussion and HIFR testing. The authors additionally thank the reviewers and editor for their thoughtful comments that improved the manuscript. This research was supported by NSF/IOS 1556396 and NSF/IOS 1754437 to MFO and DLC.

## Author Contributions

HIFR design and testing and experimental design by MKD and AND. Data collection and analysis by MKD. Writing of the manuscript and production of figures by MKD, DLC, MFO.

## Supplementary Material

**Table 1:** Materials used in building the HIFR with prices rounded to the nearest $0.25.

| Material | Manufacturer | Cost per unit | Units needed | Total Cost | Cost per chamber |
| --- | --- | --- | --- | --- | --- |
| Plexiglass | Hardware store | \$200.00 per 6ft x 4ft sheet | 1 | \$200 | \$10.00 |
| Water safe silicone glue | Hardware store | \$5.00 per tube | 1 | \$5.00 | \$0.25 |
| Plastic Weld | Hardware store | \$6.00 per 50mL | 1 | \$6.00 | \$0.30 |
| 2mm inner diameter x 4mm outer diameter tygon tubing | Amazon | \$23.00 for 15 meters | 1 | \$23.00 | \$1.15 |
| Glass chambers | Greatglas Inc. | \$10.00 per chamber | 20 | \$200.00 | \$10.00 |
| Rubber stoppers with two 4mm holes | WidgetCo | \$2.00 per stopper | 40 | \$80.00 | \$4.00 |
| 4mm outer diameter x 50mm length capillary tubes | Amazon | \$1.25 per tube | 80 | \$100.00 | \$5.00 |
| One-way valves with luer locks | Amazon | \$1.00 per valve | 60 | \$60.00 | \$3.00 |
| 0.32 cm barb male luer locks | Amazon | \$11.00 for 25 | 40 | \$22.00 | \$1.10 |
| 0.32 cm barb female luer locks | Amazon | \$11.00 for 25 | 50 | \$22.00 | \$1.10 |
| Peristaltic pumps, 12V motor, tygon tubing | Williamson Manufacturing (UK) | \$50.00 | 10 | \$500.00 | \$25.00 |
| Submersible pumps for flushing | Amazon | \$20.00 | 2 | \$40.00 | \$2.00 |
| Electronics |  |  |  |  |  |
| Arduino Uno | Amazon | \$20.00 | 1 | \$20.00 | \$1.00 |
| Double Pole Double Throw Relays | Amazon | \$10.00 for 10 | 10 | \$10.00 | \$1.00 |
| AC/DC Converter Power Supply Box | DigiKey | \$30.00 per box | 2 | \$60.00 | \$2.00 |
| Black and red electrical wires | DigiKey | \$30.00 for 50m | 2 | \$60.00 | \$2.00 |
| Outlet box | Amazon | \$5.00 for 10 | 2 | \$5.00 | \$0.25 |
| 5V relay | Amazon | \$5.00 for 2 | 1 | \$5.00 | \$0.25 |
| Replacement power cable | Amazon | \$5.00 | 1 | \$5.00 | \$0.025 |
| **Cost without Oxygen Sensing Equipment** | | | | **\$1,423.00** | **\$71.25** |
| Oxygen Sensing Technology Costs |  |  |  |  |  |
| PreSens OXY10 oxygen meter | PreSens | \$13,966.00 | 1 | \$13,966 | \$698.50 |
| PreSens flow through cells for oxygen (FTC-PSt7-10-YOP) | PreSens | \$85.00 | 10 | \$850.00 | \$42.50 |
| PreSens fiber optic cable oxygen sensors (POF-FTC-L2.5-1ST) | PreSens | \$70.00 | 10 | \$700.00 | \$35.00 |
| PreSens PT100 temperature probe | PreSens | \$163.00 | 1 | \$163.00 | \$8.25 |
| **Cost of Oxygen Sensing Equipment** | | | | **\$15,679.00** | **\$784.25** |
| **Total Cost** | | | | **\$17,102.00** | **\$855.50** |

**Figure S1.**
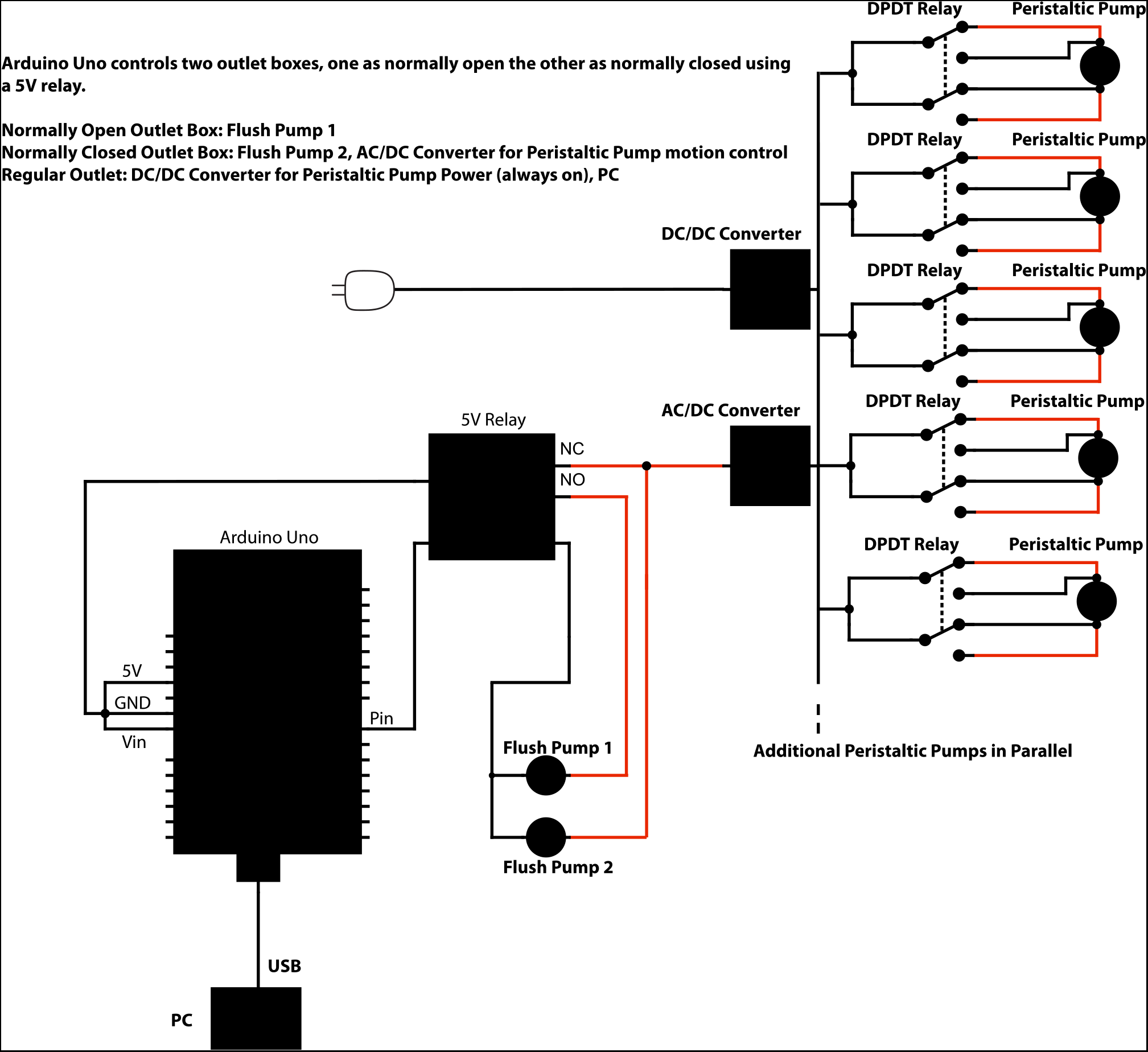
Arduino circuit and controls. Electrical circuit controlling high-throughput intermittent flow respirometer circulation and flush pumps. Note that the voltage of the power supply boxes depends on the voltage required to run the peristaltic pumps (here 12VDC) and double pole double throw relays (here 5VDC).

**Figure S2:**
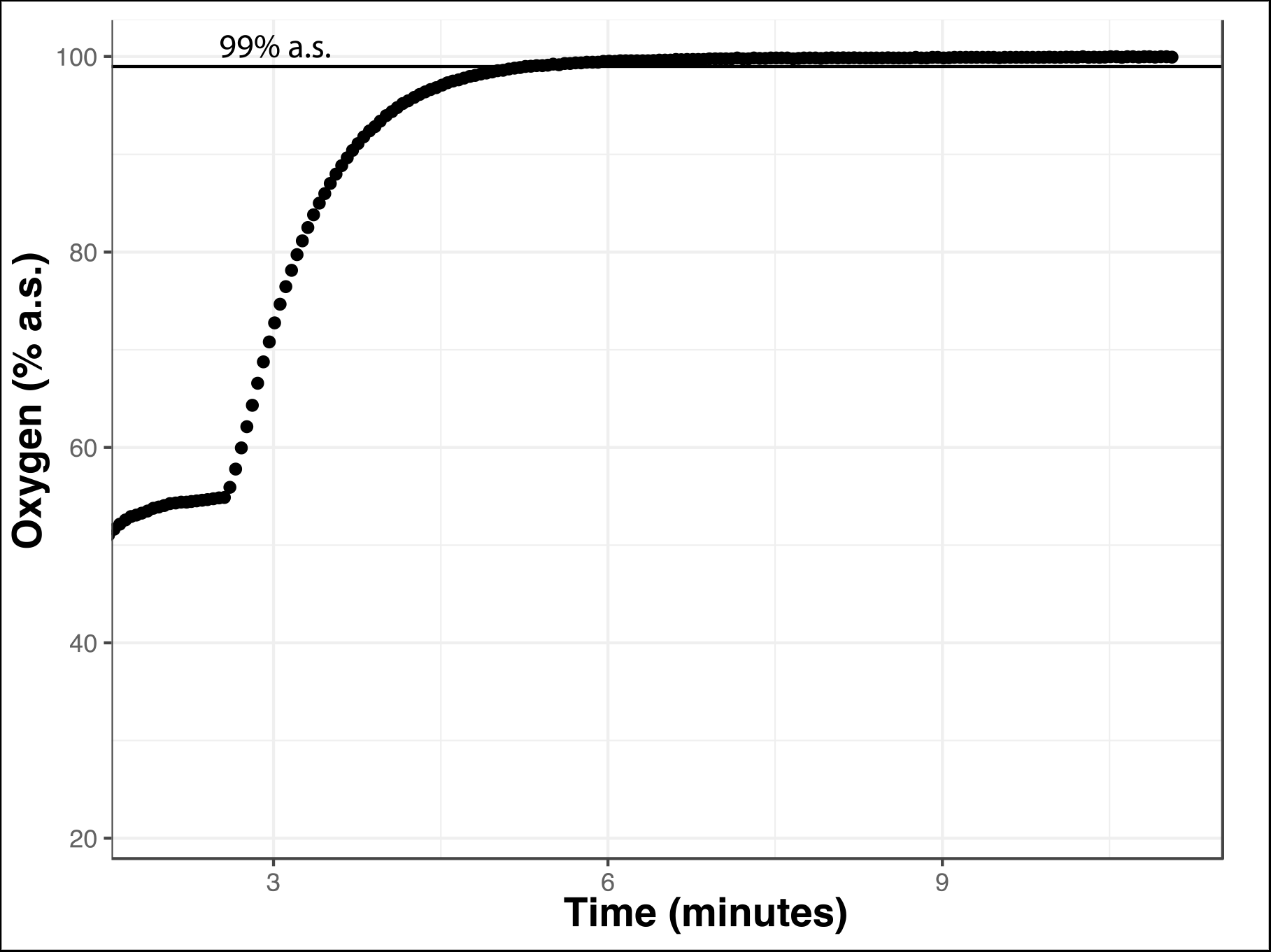
Oxygen Over Time During Flush Period. A chamber was filled with water at a low oxygen concentration (achieved by bubbling in nitrogen) and sealed before turning on the flush pump. Between 4 and 5 minutes after turning on the flush pump oxygen reached greater than 99% air saturation as expected based on steady state transformation equation (t(99%) = −ln(1 − 0.99) × 300 mL / 300 mL/min = 4.61 min, (6)).

**Figure S3:**
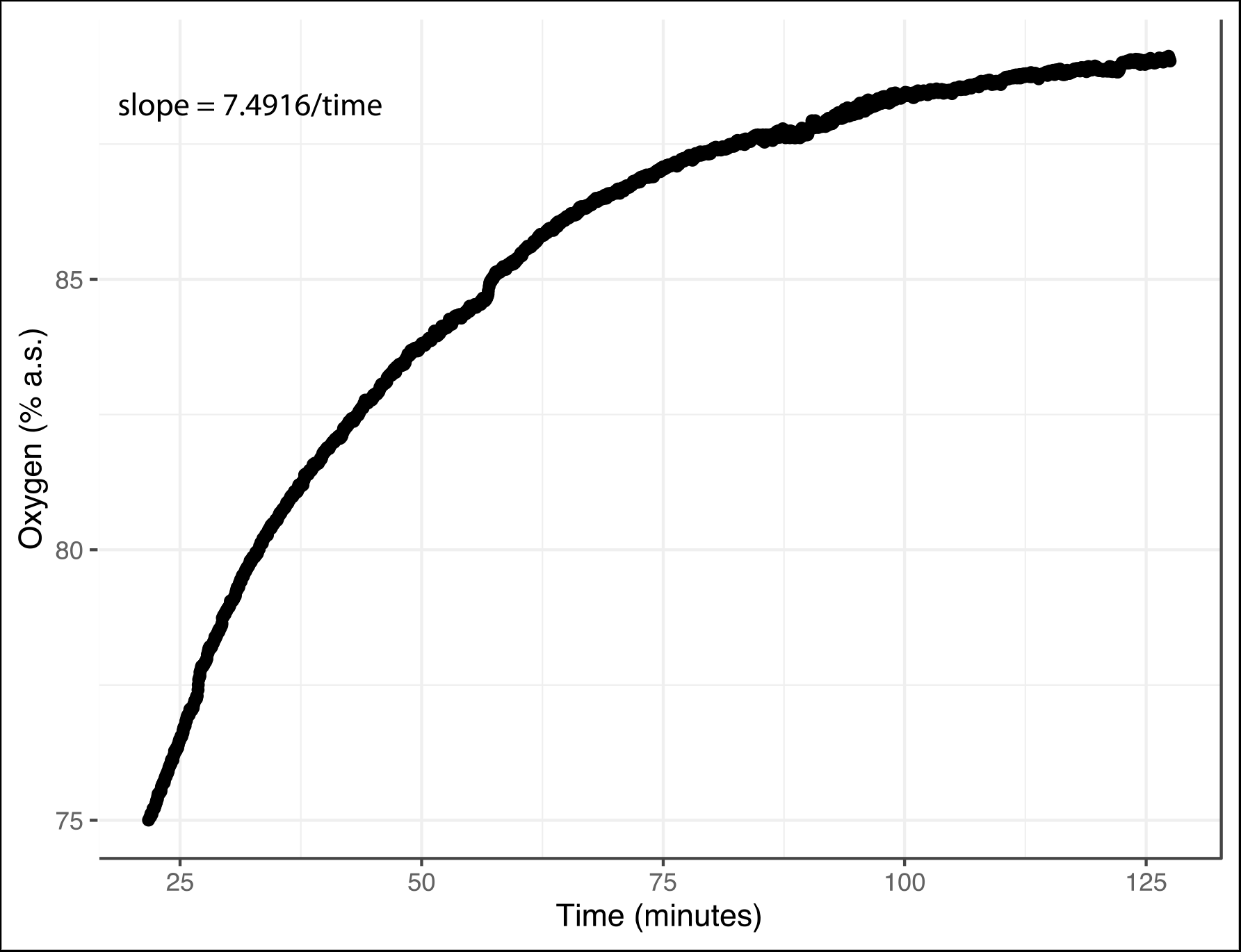
Leak of Oxygen over Time in a Sealed Chamber. A chamber was filled with water at a low oxygen concentration (achieved by bubbling in nitrogen) and sealed while the paired chamber was flushed. A model of oxygen *versus* log10(time) was used to derive an equation that can be used to predict the amount of leak at a specific time point during the test: slope (% a.s. per minute) = 7.4916/time (minutes). Leak did not exceed 0.14% a.s. per minute from 85 to 89% a.s. and decreased as the oxygen concentration in the chamber increased.

